# Novel 4,400-year-old ancestral component in a tribe speaking a Dravidian language

**DOI:** 10.1101/2024.03.31.587466

**Authors:** Jaison Jeevan Sequeira, Swathy Krishna, George van Driem, Mohammed Shafiul Mustak, Ranajit Das

**Affiliations:** Yenepoya Research Centre, Yenepoya (deemed-to-be) University, Mangalore, Karnāṭaka, India; Department of Applied Zoology, Mangalore University, Mangalagangothri, Karnāṭaka, India; Institut für Sprachwissenschaft, Universität Bern, Länggassstrasse, Bern, Switzerland

**Keywords:** Koraga, North Dravidian, Dravidian, Elamo-Dravidian, Indus Valley, Harappan civilisation, ANI, ASI

## Abstract

Research has shown that the present-day population on the Indian subcontinent derives its ancestry from at least three components identified with pre-Indo-Iranian agriculturalists once inhabiting the Iranian plateau, pastoralists originating from the Pontic-Caspian steppe and ancient hunter-gatherer related to the Andamanese Islanders. The present-day Indian gene pool represents a gradient of mixtures from these three sources. However, with more sequences of ancient and modern genomes and fine structure analyses, we can expect a more complex picture of ancestry to emerge. In this study, we focus on Dravidian linguistic groups to propose a fourth putative source which may have branched out from the basal Middle Eastern component that gave rise to the Iranian plateau farmer related ancestry. The Elamo-Dravidian theory and the linguistic phylogeny of the Dravidian family tree provide chronological fits for the genetic findings presented here. Our findings show a correlation between the linguistic and genetic lineages in language communities speaking Dravidian languages when they are modelled together. We suggest that this source, which we shall call ‘Proto-Dravidian’ ancestry, emerged around the dawn of the Indus Valley civilisation. This ancestry is distinct from all other sources described so far, and its plausible origin not later than 4,400 years ago on the region between the Iranian plateau and the Indus valley supports a Dravidian heartland before the arrival of Indo-European languages on the Indian subcontinent. Admixture analysis shows that this Proto-Dravidian ancestry is still carried by most modern inhabitants of the Indian subcontinent other than the tribal populations. This momentous finding underscores the importance of population-specific fine structure studies. We also recommend informed sampling strategies for biobanks and to avoid oversimplification of ancestral reconstruction. Achieving this requires interdisciplinary collaboration.

## Introduction

Numerous studies have highlighted the genetic complexity of modern Indian populations, which include primitive hunter-gatherer tribes, dry and wet land agriculturist communities, pastoralists, warrior clans, trading communities, artisans and priestly castes ^1–5^. In the last two decades, we have seen a transition from SNP chip technology to whole genome sequencing. Currently, with the increased number of ancient and modern genomes, population genetics has been transformed into an interdisciplinary enterprise involving archaeologists, historians, linguists and ethnographers, thereby inviting public attention and debate around complex questions of our past. The genetic source of present-day Indians remains a topic of utmost interest. Numerous studies have used genotypic information to reveal the complex ancestral components in the gene pool of the Indian subcontinent. Reich et al. (2009) ^6^ proposed two broad ancestral components, i.e. Ancestral North Indians and Ancestral South Indians, with varying proportions of Andamanese ancestry and steppe pastoralist ancestry. Subsequently, attempts were made to understand the correlation of these ancestral components with language families and geography ^3^. The most recent work involving 2,700 Indian genomes suggests three putative ancestral sources for modern Indians that have been labelled ‘Iranian plateau farmer related’, ‘Pontic-Caspian steppe pastoralist related’ and ‘Andamanese hunter-gatherer related’ ^7^, but we suggest that this is an oversimplification.

Interdisciplinary studies investigating local ancestry using adaptive markers, linguistic affinity, cultural similarities and social affiliation testify to the complexity of genetic structure in present-day populations. A recent study suggests that the U1 macrohaplogroup in the Koraga tribe could be a correlate of Dravidian linguistic lineage ^8^.

The Koraga are an indigenous tribal community who reside on the southwestern coast of India, primarily in Dakṣiṇa Kannaḍa and Uḍupi districts of Karnāṭaka and Kāsaragoḍ district of Kerala. They represent one of the most marginalised and impoverished populations in southern Karnāṭaka, and their livelihood mainly depends on activities such as basket weaving, collecting firewood and honey from nearby forests, and working as seasonal daily wage labourers. Although the Koraga language has been influenced for centuries by surrounding Tuḷu speakers, and many Koraga are bilingual in Tuḷu, Bhat ^9^ (1971) and McAlpin ^10^ (1981) grouped Koraga together with Kurukh and Malto under the North Dravidian branch. North Dravidian language communities are mainly distributed geographically across the northwest and north of the Indian subcontinent. Krishnamurti ^11–13^ classified the Brahui language, spoken in Belochistan in the far northwest, as North Dravidian, whereas McAlpin ^10^ (1981) and Zvelebil ^14^ (1990) placed Brahui under a separate node as a distinct branch of its own within Dravidian or Elamo-Dravidian, intermediate between Elamite and mainstream Dravidian. Zvelebil ^14^ (1990) likewise proposed to treat Koraga as an independent branch of Dravidian under its own node in the tree. So, whereas these languages are sometimes lumped together under the label Northern Dravidian *sensu lato*, linguists have also seen Brahui, Northern Dravidian *strictu sensu* and Koraga as possibly representing three primary branches of the language family. By contrast, a fourth branch, dubbed ‘mainstream Dravidian’ by McAlpin ^10^ (1981), encompasses most modern Dravidian languages and consists of a southern subgroup (i.e. Tamiḻ, Malayāḷam, Iruḷa, Tōḍa, Kota, Koḍagu, Kuṟumba, Baḍaga, Kannaḍa, Tuḷu), a south-central subgroup (i.e. Telugu, Gondi, Koṇḍa, Maṇḍa, Pengo, Kuvi, Kui) and a central subgroup (i.e. Kolami, Naikri, Naiki, Gadaba, Parji). The objective of this study was to determine the ancestral origin of the Koraga tribe in this context and therefore the roots of the Dravidian family. In order to achieve this, we utilised single nucleotide polymorphisms (SNP). We genotyped 29 unrelated Koraga individuals using the Infinium Global Screening Array-24 v3.0 (GSA v3.0) BeadChip platform and utilised population genetic tools to trace their ancestry.

## Results

### Population structure in the Indian cline

We performed PCA analysis to observe the position of Koraga in the Indian cline. Koraga samples clustered with Gauḍa, Bāgdi and Tānti samples (Figure 1 a). Geographically these castes are found in the northern and eastern zones. This cluster appears to have drifted away from the Ancestral North Indian to Ancestral South Indian cline, parallel to the southwestern and eastern Indian clusters. Interestingly, the clusters with North Dravidian speaking tribes at the Ancestral South Indian end appeared closer to each other, whereas the Brahui samples clustered at the Ancestral North Indian end. The geographical locus of the primary branches of the Dravidian language family lies in the northwest, where the Brahui language community has survived as a remnant population, whereas today most modern Dravidian language speakers live in South India ^14,15^. It is evident from the PCA plot that these groups have drifted away with time (Figure 1 c).

**Figure 1.**
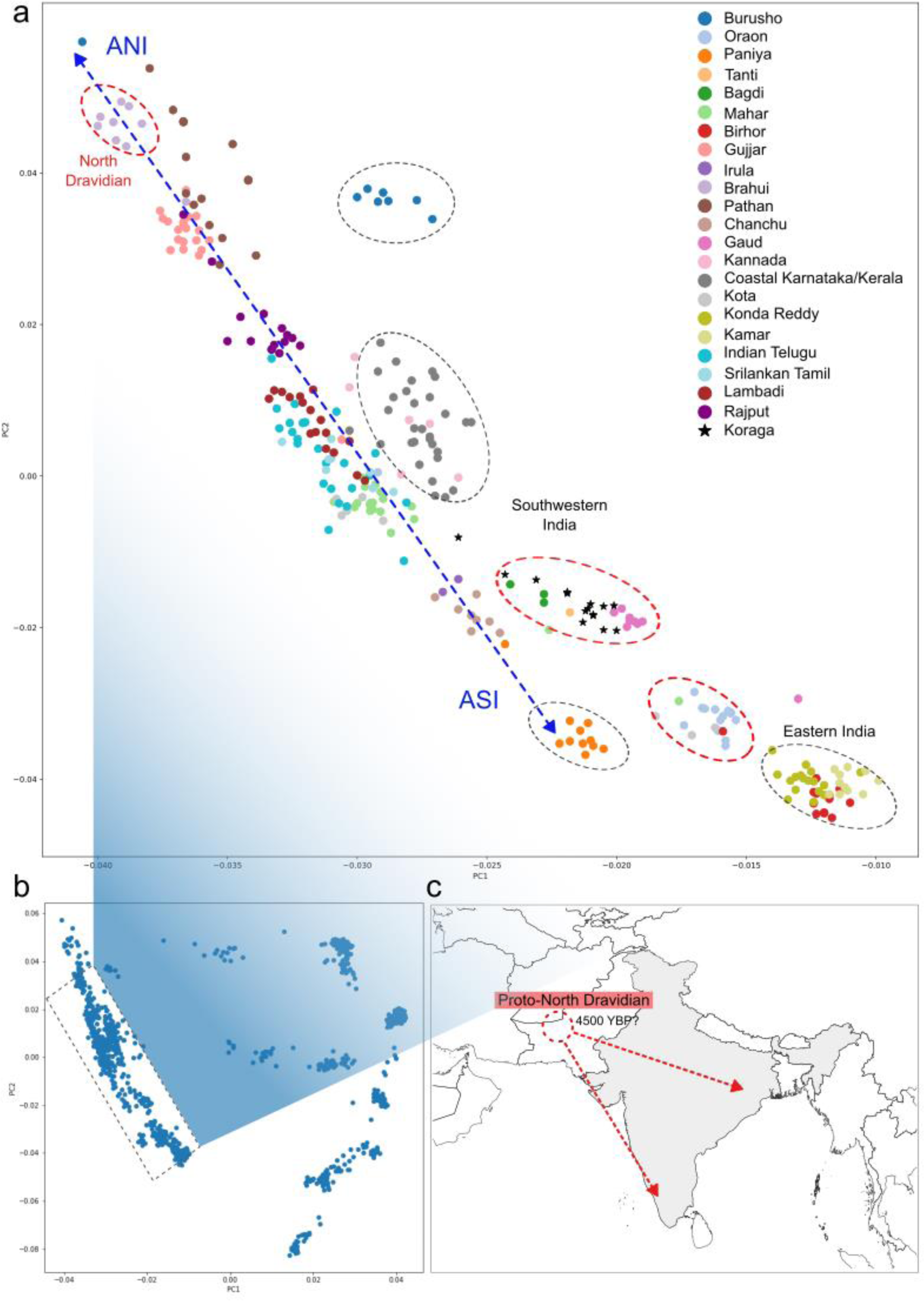
Population structure in the Indian cline. (a) PCA plot show a clustering pattern along the ANI-ASI cline. Red ellipses represent clusters which also include North Dravidian language communities sensu lato: Brahui, Koraga, Gauḍa, Tānti, Bāgdi, Oraon, Kota, Birhor and Mahar. Kota and Mahar are found in the ANI-ASI cline as well. Grey ellipses represent clusters that have drifted away from the ANI-ASI cline, which includes both Dravidian and other groups: Burusho, Kannaḍa and coastal populations of Karnāṭaka and Kerala, Paniya, Koṇḍa Reddy, Kamar and one Birhor sample. (b) Overall PCA plot for GenomeAsia100K including Koraga samples. (c) Putative migration routes for Dravidian tribes.

### Admixture profile and ancestry of the Koraga tribe

In order to estimate admixture dates between the geographically separated tribes speaking Dravidian languages, we performed ALDER analysis. This test computes LD decay curves and performs curve-fitting to estimate the admixture dates. Admixture between major North Dravidian tribes *sensu lato*, viz. Koraga, Brahui and Oraon (Kurukh), happened between 5,988 and 2,800 years ago (Table 1). The median falls around 2,370 BC, coinciding with the Mature Harappan period. Notably, the Oraon and Birhor populations are presently found in eastern India, and Brahui is found in Belochistan. Linguistically, both the Koraga and Brahui languages belong to the North Dravidian lineage, whilst North Dravidian language communities are found in eastern India as well. Our findings substantiate the hypothesis of a Dravidian homeland before the arrival of Indo-European languages into the Indian subcontinent.

**Table 1.**
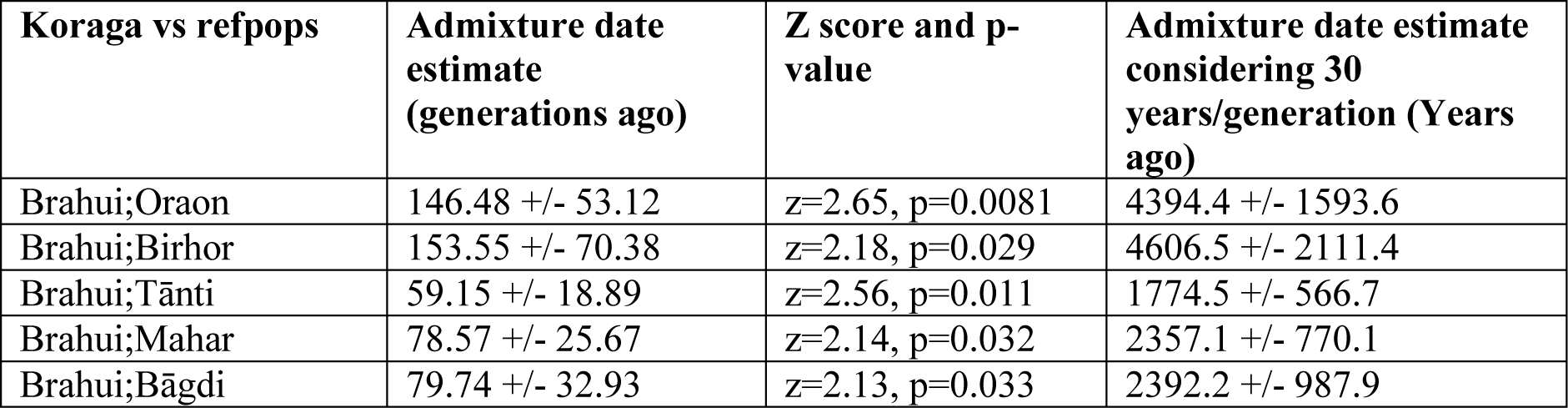
Date estimates for admixture between Koraga, Brahui and other populations.

Furthermore, to confirm admixture events between the North Dravidian tribes, Admixtools 2 produced the best fit when Koraga, Brahui and Oraon shared ancestry (Supplementary Figure 3). Admixture graphs are more meaningful when models include ancient and modern samples. Therefore, we repeated the same analysis using Dataset 2 containing 4605 modern and ancient samples. In spite of a lesser number of SNPs, the admixture graph produced the model which fit best, at a score of 2.97, when the ancestral Koraga diverged from the Andamanese ancestor carrying 66% of this ancestry (Figure 2 b). The remaining 34% was derived from the ancestor of the 10,000-year-old Neolithic sample from Ganj Dareh, in the Zagros mountains of what today is western Iran. Notably, the Indus Periphery samples did not share any ancestry with the Koraga tribe (Fig 2 a, b, c, e). “Indus Periphery” is the label given to ancient DNA samples dating from the 4th–3rd millennium BC from Gonur in what today is Turkmenistan and Shahr-i-Sokhtah in what today is eastern Iran. To corroborate this key finding, we included Paniya, a hunter-gatherer tribe, and Kuruba, a pastoralist tribe, in the model. Whilst Paniya formed a separate branch (Figure 2 e), the pastoralist tribe Kuruba turn out to have received their ancestral components from an Andamanese-ancestor-related population and the ancestor of the Indus Periphery samples (Figure 2 c, d, e). Koraga, on the other hand, remains unrelated to the Indus Periphery, although at a deeper level the Koraga ancestor shared 39% of his ancestry with the Indus Periphery lineage (Figure 2 d).

**Figure 2.**
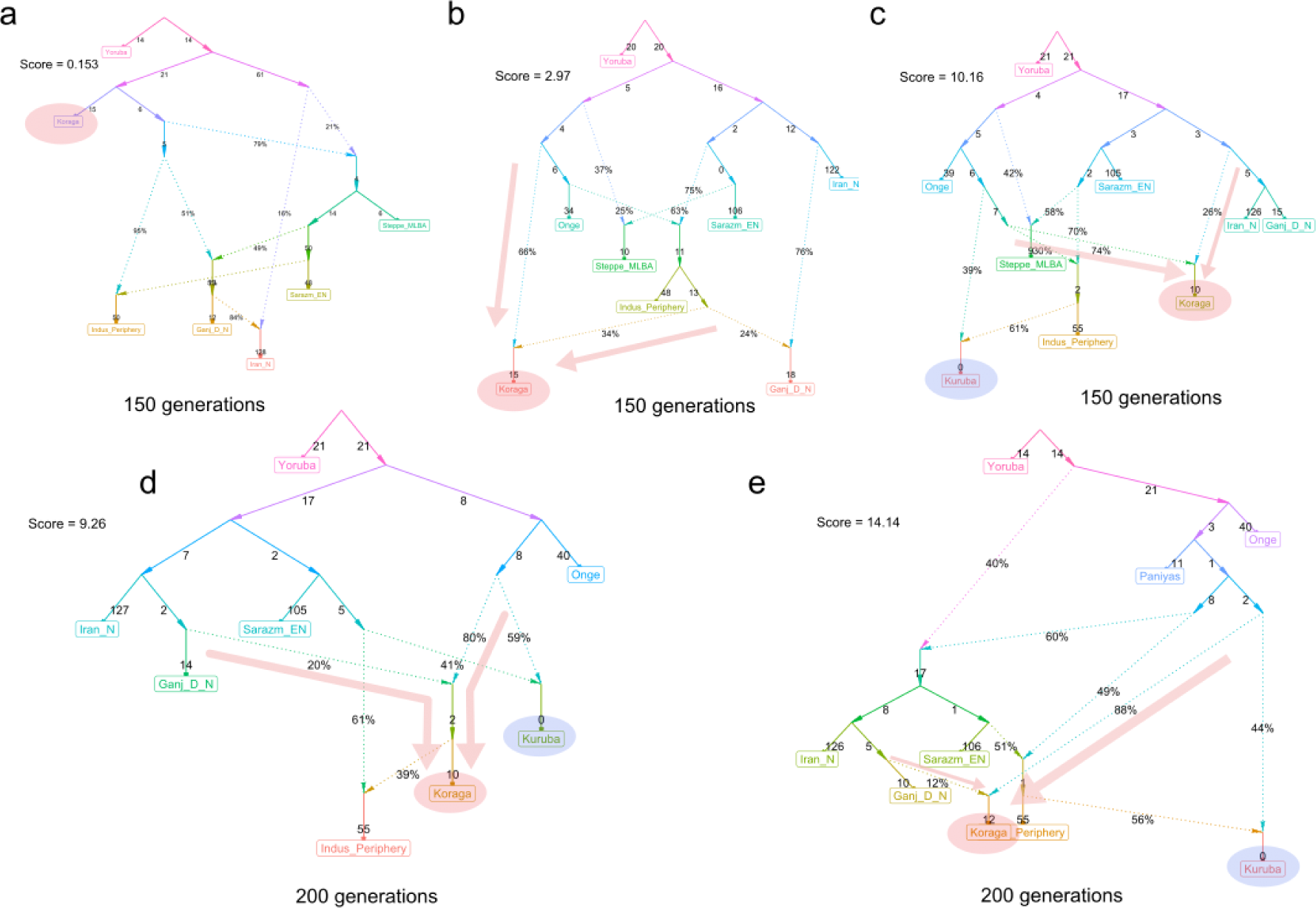
Admixture graphs show different models for Koraga ancestry. Koraga is represented in the red ellipse, and Kuruba is represented in the blue ellipse. Arrows show the most recent admixture event for the Koraga tribe. N=Neolithic, EN=Eneolithic.

We used f3 statistics to distinguish between known ancestries and to compare them with the Koraga ancestry. We used Hàn and Yoruba as outgroups. A significant negative z-score indicates that admixture has taken place between the three populations tested. It has been established that Önge and the “Indus Periphery” represent two of the major ancestral components in populations of the Indian subcontinent^16^. So, we compared their significant z-scores with those of Koraga population. Notably, the significant z-scores, for which higher values indicate a lower probability of admixture, are comparable between Önge, Indus Periphery and Koraga (Supplementary Table S1). However, the genetic distance between Koraga (∼0.015) and other populations is not as great as between Önge (∼0.041) and the Indus Periphery (∼0.060) and other ancient and modern populations.

The statistical tool qpAdm, implemented in Admix Tools v5.1 ^17^, was employed to estimate the ancestry proportions in the Koraga genomes originating from a mixture of reference populations. The Koraga were modeled as a combination of three source populations, namely Önge, Indus Periphery and a third population from Eurasia and Africa (See Materials and Methods). We found that the Koraga can be best modeled as the genomic admixture of Önge, the Indus periphery and modern-day Iranians (Table 2). Whilst discernible ancestry fractions from various countries across the Middle East were identified in the Koraga, the Mid to Late Bronze Age steppe ancestry (MLBA) and African (Yoruba) ancestry fractions were found to be the least prominent (Table 2). This finding further refutes any possibility of an African origin for the Koraga. Our findings align with the recent mitochondrial DNA results ^8^, affirming that the maternal ancestry of the Koraga can be traced back to the Middle East, with discernible fractions of West Eurasian ancestry in their genomes.

**Table 2:**
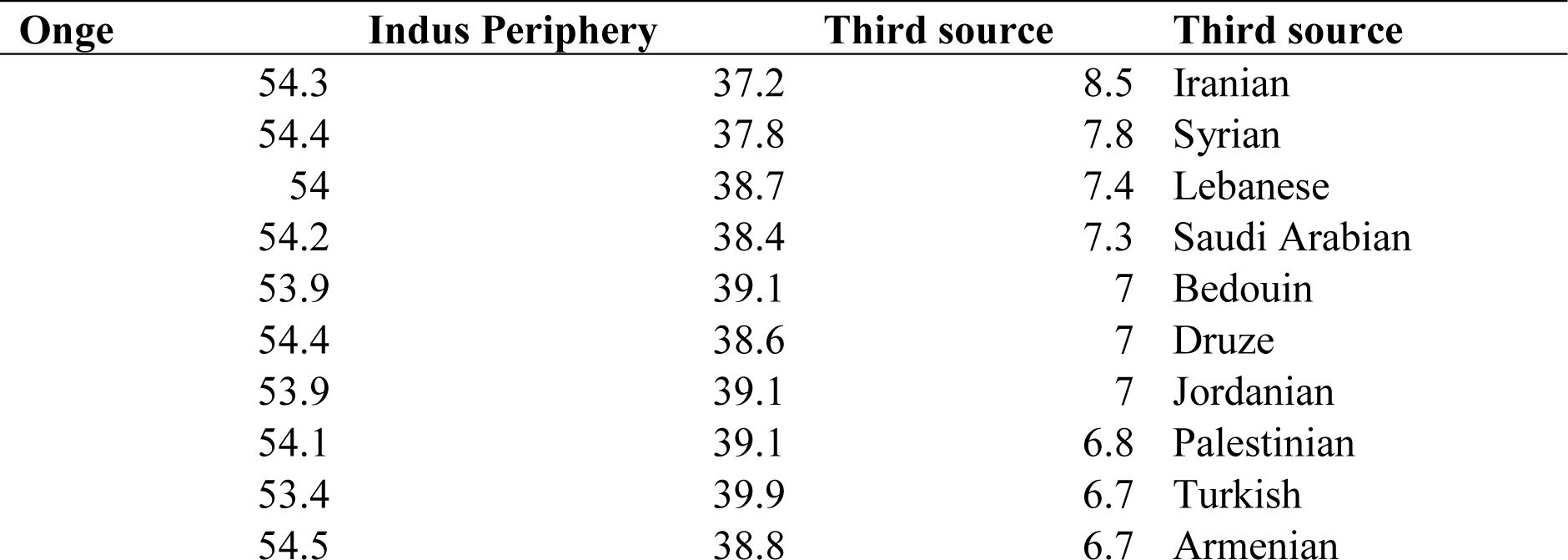
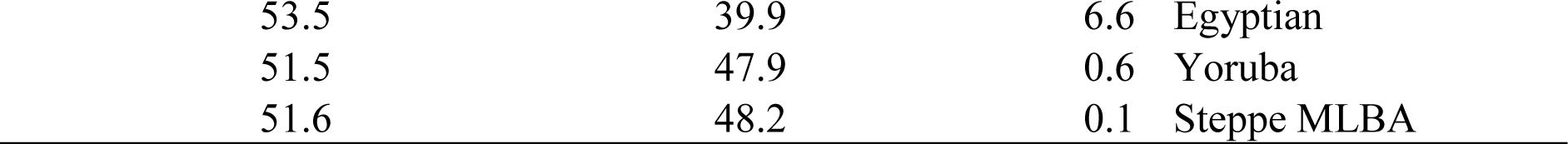
Ancestry fractions (%) in Koraga genomes. The Koraga were modeled as a combination of three source populations, namely Önge, the Indus periphery and a third population from Eurasia and Africa.

| Önge | Indus Periphery | Third source | Third source |
| --- | --- | --- | --- |
| 54.3 | 37.2 | 8.5 | Iranian |
| 54.4 | 37.8 | 7.8 | Syrian |
| 54 | 38.7 | 7.4 | Lebanese |
| 54.2 | 38.4 | 7.3 | Saudi Arabian |
| 53.9 | 39.1 | 7 | Bedouin |
| 54.4 | 38.6 | 7 | Druze |
| 53.9 | 39.1 | 7 | Jordanian |
| 54.1 | 39.1 | 6.8 | Palestinian |
| 53.4 | 39.9 | 6.7 | Turkish |
| 54.5 | 38.8 | 6.7 | Armenian |
| 53.5 | 39.9 | 6.6 | Egyptian |
| 51.5 | 47.9 | 0.6 | Yoruba |
| 51.6 | 48.2 | 0.1 | Steppe MLBA |

We then performed a model-based population structure analysis, using ADMIXTURE v1.3. Dataset 1 was used for this purpose, as this set covered 5,53,102 SNPs, thus providing greater reliability. The lowest cross-validation error (CVE) was observed for K = 11 (Supplemental Figure 2). At K = 11, Indonesians (K1, orange), East Asians (K2, yellow), Filipinos (K3, green), Papuans (K4, navy blue), Buryats (K5, purple), the Jarawa and Önge (K6, red), the Tōḍa from Tamiḻ Nāḍu (K7, light green), Koraga (K8, cyan), British (K10, violet) and Yorubans from Africa (K11, dark green) were assigned to distinct clusters (Figure 3). The tribal populations exhibited higher proportions of an indigenous Indian component (K9, blue), along with moderate proportions of K1 and minor fractions of K6, indicating their ancestral origin associated with these two populations. Consistent with prior studies ^2,3,18,19^, the admixture plot revealed that mainstream Indian populations were comprised of variable proportions of light green, cyan and blue (K7, K8 and K9, indigenous Indian components, likely ASI-related), violet (K10, West-Eurasian ancestry fraction), and minor fractions of red (K6, likely derived from Ancient Ancestral South Indians: AASI populations) and yellow (K2, East Asian ancestry fractions). In particular, the Manipuri genome employed in this plot revealed a high fraction of yellow (K2) with minor fractions of K8, K9 and K10, potentially linked to a common origin and admixture history.

**Figure 3.**
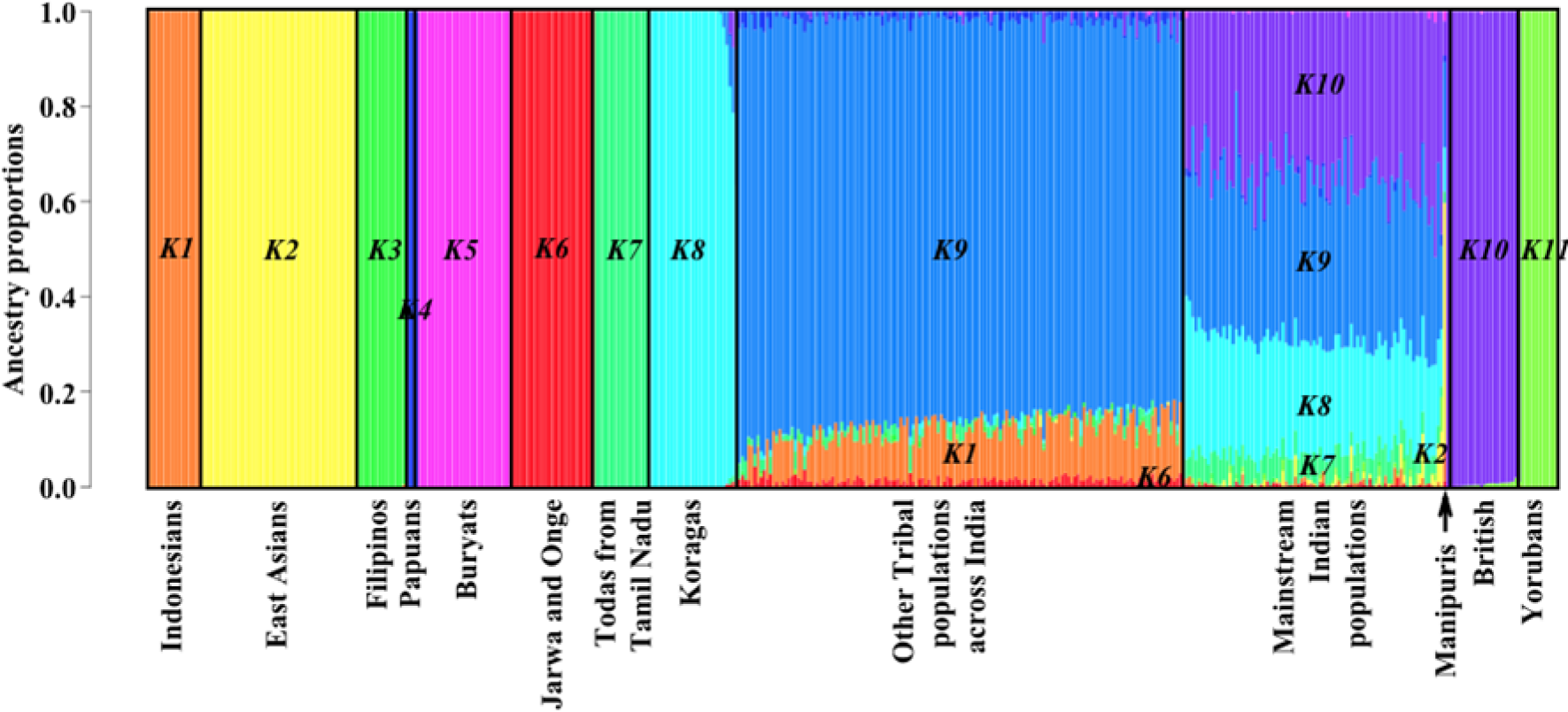
Admixture analysis of 1,279 individuals from Dataset 1. The admixture plot illustrates the ancestry components of the samples across the globe. Admixture proportions were obtained through unsupervised analysis at K = 11, using ADMIXTURE v1.3 and visualised in R v3.5.1. Each individual is depicted by a vertical line, segmented into coloured sections. The lengths of these segments are proportionate to the contributions of ancestral components to an individual’s genome.

**Figure 4.**
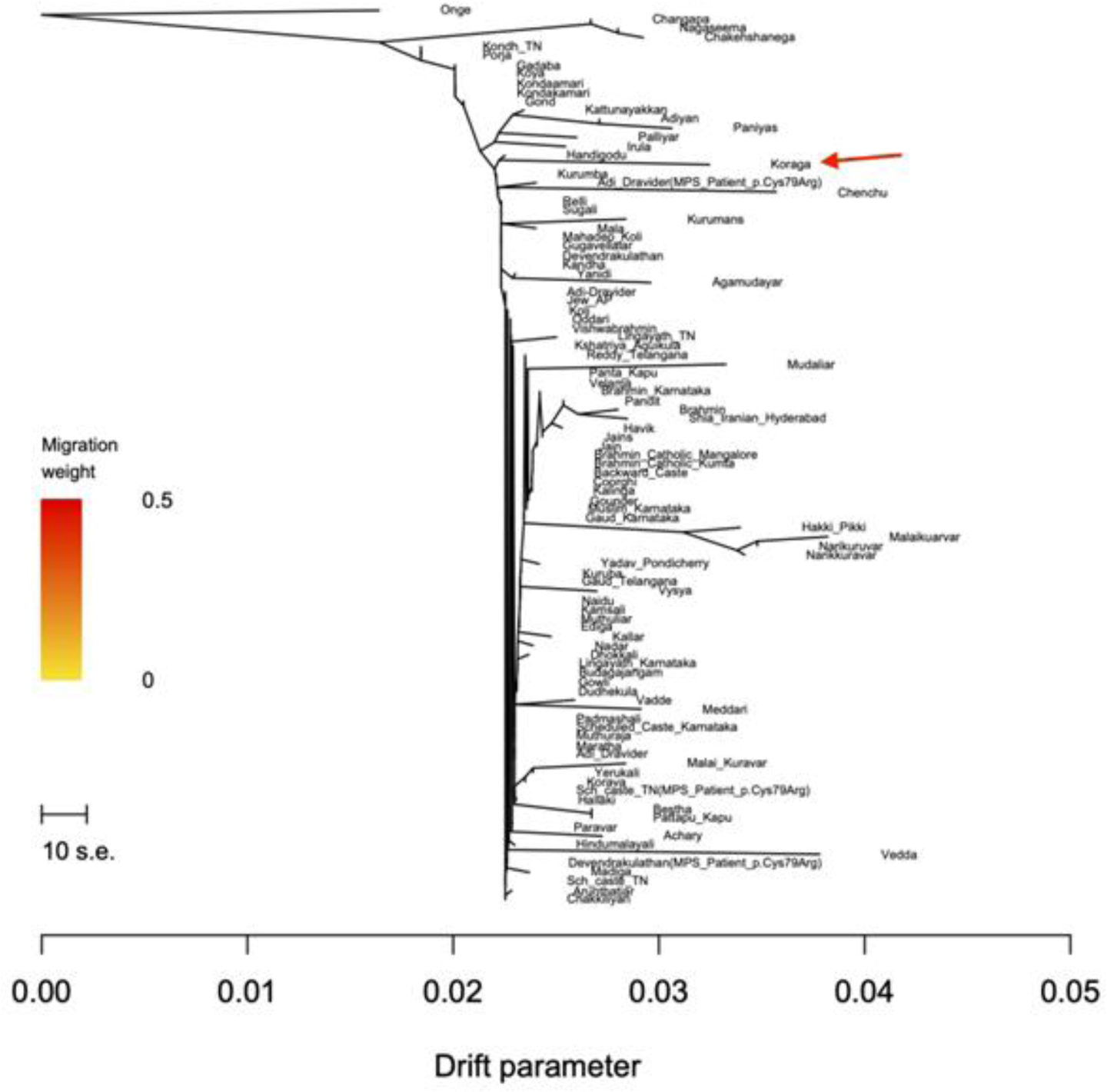
Maximum Likelihood (ML) tree showing the genetic relatedness between the Koraga and selected South Asian populations present in Dataset 3, using TreeMix v1.13. The tree was rooted with Önge, an Andamanese population. The horizontal axis represents the drift parameter, and the scale bar indicates ten times the average standard error of the entries in the sample covariance matrix. The ML tree depicted notable genetic relatedness between the Koraga and Handigōḍu people of Karnāṭaka. Additionally, the ML tree highlighted their genetic similarities with various indigenous South Indian populations, including Paniya, Iruḷa and Kuruba.

Except for three samples, the majority of Koraga genomes analysed in this study do not exhibit K9 and K10, suggesting an early divergence from mainstream Indian populations. This finding supports our hypothesis that Koraga ancestry diverged from other ancestries found on the Indian subcontinent. Moreover, the complete absence of the K11 component reflects the likely absence of ancestral links between the Koraga and African populations.

We utilised TreeMix v.1.13 to explore the patterns of population splits and admixtures amongst selected South Asian populations, using Dataset 3. The maximum likelihood (ML) tree generated by TreeMix using Dataset 1, revealed a high degree of genetic relatedness between the Koraga and Handigōḍu people from the Śivamogga (Shimoga) district of Karnāṭaka. The Handigōḍu, much like the Koraga, are a small isolated South Indian population with a high prevalence of Handigōḍu syndrome, a rare autosomal dominant form of spondyloepimetaphyseal dysplasia (PMID: 7886470). Similar to the Koraga, many Handigōḍu die at a very early age (25-30 years). The ML tree also indicated a genetic affinity of the Koraga with other primitive indigenous South Indian groups, including the Paniya, Iruḷa, Kuruba, and Adi Drāviḍa from Tamiḻ Nāḍu.

### Founder event in the Koraga tribe

Previous studies have revealed that many Indian tribes exhibit high levels of founder effect and inbreeding ^20,21^. We utilised ASCEND software to determine the date of the founding event of the Koraga tribe. Our analysis revealed that the most recent founder event occurred between 750 and 1020 years ago, with an intensity of 6.7% to 8.2%. This intensity is approximately five times stronger than that observed in the Ashkenazi Jews ^20^. During this time, the Kadamba dynasty ruled the region, but their reign was disrupted by the rule of the Rāṣṭrakūṭa, Hoysaḷa and Western Cālukya dynasties, whilst the Perumāḷ dynasty ruled in Kerala to the south. From temple records and inscriptions, it is evident that Brahmins, who had settled along the southwestern coast, had begun to exert significant influence under the patronage of local rulers ^22^. Imposition of Brahmanism and more rigid observance of the caste system may have increased social exclusion due to the practice of untouchability, which led to the genetic isolation of the tribal populations. These developments may have contributed to the gradual reduction of the Koraga population and left genetic consequences. Our research, along with the findings of a recent study on the maternal ancestry ^8^, indicates that the formation of the Koraga gene pool began not later than 2,000 years ago, followed by a founder event about 1,000 years ago. Consequently, this intact gene pool can serve as an appropriate proxy in the absence of ancient DNA in southern India.

## Discussion

Earlier studies on the Koraga tribe using uniparental markers revealed unusually high frequencies of the U1 and H1 haplogroups in the maternal and paternal ancestry respectively ^8,23^. This finding was considered unusual because the maternal U1 haplogroup is West Asian, whereas the paternal H1a is Indian-specific. Such a contrast is either the result of a complete turnover or a bottleneck event. Sequeira et al. 2024 concluded that the U1 haplogroup is a correlate for the Koraga language, which belongs to the North Dravidian branch *sensu lato* of the Dravidian language family. Against this backdrop, we have now investigated the ancestral origin of the Koraga tribe using autosomal SNPs.

The exceptional nature of the Koraga gene pool was evident in the overall F_st_ as well. The Koraga clustered away from most of the Northern and Southern populations. Whereas in the PCA plot, Koraga samples scattered in a pattern similar to that of another North Dravidian speaking tribe from the eastern frontier. Both these tribes clustered away from Brahui, a North Dravidian speaking remnant population that through genetic assimilation now clusters with Indo-European speaking groups. The geographical distribution of North Dravidian language communities has been pointed out by linguists for over 200 years. Our findings are an attempt to understand the geographical range of North Dravidian based on admixture models and dating. Using ALDER admixture date estimation, we demonstrate that the three major North Dravidian speaking populations shared a common ancestor about 4,500 years ago. Both f3-statistics and admixture graphs show that the Brahui and Oraon (Kurukh) underwent different demographic changes as compared with the Koraga tribe, whose gene pool remained relatively intact until experiencing a strong bottleneck 1,000 years ago, possibly as a result of social exclusion. Ancestral connections between the Koraga and the 10,000-year-old Neolithic sample from Ganj Dareh in the Zagros mountains of eastern Iran points to a shared ancestry at the putative time depth ascribed by linguists to Elamo-Dravidian, a proto-language ancestral to both the Elamite language and the Proto-Dravidian language hypothetically spoken by the inhabitants of the Mehrgarh Neolithic that represented the direct antecedents to the Indus Valley civilization ^5,15,24,25^. Both the Koraga and the Early Neolithic Ganj Dareh sample share a common ancestor, and we observe the Koraga component (K8 in the ADMIXTURE bar plot) in most of the present-day non-tribal populations alongside and in addition to the Iranian plateau farmer related, Pontic-Caspian steppe pastoralist related and Andamanese hunter-gatherer related components. The bias in the contribution of a West Asian maternal ancestry to the Koraga gene pool and its correlation with the Koraga mother tongue, as well as the geographical spread of Dravidian languages allows us to hypothetically identify this novel ancestry tentatively as Proto-Dravidian. The descendants bearing this ancestry dispersed throughout the Indian subcontinent as pastoralists and farmers and gave rise to the populations that formed today’s Dravidian language communities. In this context, the question must be addressed as to what relationship obtains with the population that peopled the Indus Valley civilisation.

A recent study suggests that archaic DNA (Sarazm_EN) dating from the 4th millennium BC from what today is Tajikistan as the best proxy for Iranian plateau farmer-related ancestry ^7^. We do not find any direct ancestry sharing between our hypothetical Proto-Dravidian and Sarazm_EN or with the Indus Periphery cline. However, the f3-statistics for our hypothetical Proto-Dravidian are similar to those of Sarazm_EN, the Indus Periphery cline as well as the ancient DNA dating from 9th–8th millennium BC in the Zagros mountains, i.e. Iran_N and Ganj Dareh_N, suggesting a pre-Neolithic common ancestor related to the ancient Caucasus hunter-gatherer component that diverged from the Andamanese hunter-gatherer lineage in the Late Pleistocene ^26^. Our putative Proto-Dravidian ancestry therefore evidently constituted a separate entity that existed alongside the Iranian plateau farmer related ancestry since the Neolithic period through the Chalcolithic in the vicinity of Indus Valley civilisation. The Elamo-Dravidian theory and the linguistic phylogeny of the Dravidian family tree provide ideal chronological fits for the genetic findings presented here. The time depth of the shared ancestry between the Koraga and Early Neolithic Ganj Dareh 10,000 years ago coincides with the time ascribed by linguists to the hypothetical Elamo-Dravidian linguistic phylum in the Early Holocene and matches geographically with the Elamo-Dravidian homeland in the Zagros mountains, as proposed by McAlpin ^10^ (1981). The dating of the ‘Proto-Dravidian’ ancestry component matches the flourishing of the Indus Valley civilisation before the demise and break-up of Harappan civilisation. Subsequently, this ancestral component diffused throughout the Indian subcontinent except into tribal populations, whose indigenous ancestry in the Indian subcontinent antedates the time of the Dravidian diffusion.

## Materials and Methods

### Sample collection

Saliva samples were collected in compliance with the institutional ethical guidelines of the Yenepoya Ethical Committee 1 (YEC-1/2021/052), affiliated with the Yenepoya deemed-to-be University in Mangalore with written informed consent from 29 unrelated individuals from the Koraga community in the Beḷtaṅgaḍi and Mangalore taluks of Dakṣiṇa Kannaḍa district in Karnāṭaka. DNA was extracted by the non-invasive MagStable DNA Saliva Collection Kit (MagGenome Technologies Pvt. Ltd., Cochin). Genotyping was performed using the Infinium Global Screening Array-24 v3.0 (GSA v3.0) BeadChip Illumina Inc, California. This platform comprised 648,465 Single Nucleotide Polymorphism (SNP) markers. The study participants included 17 males and 12 females, specifically from the Kuṇṭu (N=7) and Tappu (N=22) Koraga clans, ranging from 20 to 70 years of age.

### Data curation and dataset generation

Four datasets were created for downstream analysis. In Dataset 1 (N=1,279), the genotype data from 29 Koraga samples were merged with 87 non-Koraga Indian samples, genotyped on the GSA v3.0 platform, available in our in-house database and 1,163 samples from the GenomeAsia100K Consortium (PMID: 31802016), assessing 5,53,102 SNPs. For Dataset 2 (N=4,605), the genotype data from 29 Koraga samples were merged with 4,576 ancient and modern genomes across the globe, including the recently published Harappan genome from Rākhīgaḍhī, assessing 14,539 SNPs ^2,16,21,27,28^. Furthermore, a subset of Dataset 2 was generated (Dataset 3), comprised of 29 Koraga genomes and 693 other South Asian genomes, assessing 14,539 SNPs. File conversions and manipulations were performed using VCFtools v.0.1.13 ^29^, PLINK v1.9 ^30^ and EIGENSOFT v7.2.1 ^31,32^. Dataset 4 included 313 individuals from GenomeAsia100K, covering 190515 SNPs.

### Population structure and admixture analysis

To examine the population structure of the Koraga tribe and their relationship to other populations, we conducted Principal Component Analysis (PCA) using smartpca from the EIGENSOFT package (version 18140). We analysed Dataset 1, which included 1,279 individuals and 553,102 SNPs. The top two principal components were plotted using in-house python script.

To model an admixture graph, we used Dataset 2 with 12,379 SNPs from modern and ancient populations that were pruned for LD (–indep-pairwise 50 10 0.1) and fed it into ADMIXTOOLS 2 ^33^. We began by calculating pairwise f2 statistics between the groups using the “extract_f2” function. Then, we used “f2_from_precomp” to extract allele frequency products from the computed f2 blocks. For each scenario in which we were interested, we sought the best-fitting admixture graph using “find_graphs”. We selected the graph with the lowest score. First, we started with no migrations, and then gradually added migrations until we found the best-fitting graph for that scenario. We tested models for up to 200 generations.

In order to examine the genetic relationship and potential for admixture between the Koraga population and other modern and ancient populations, we utilised the f3 function in the ADMIXTOOLS 2 package within R to perform outgroup f3-statistics. Negative f3-statistics indicate admixture within the test population, whilst positive statistics suggest an unadmixed population. We designated the Yoruba and Hàn as our outgroups.

The genetic ancestry of all individuals was determined utilising an unsupervised clustering algorithm, ADMIXTURE v1.3 ^34^. We used Dataset 2 for this purpose. The optimal number of ancestral components (K) was identified by minimising the cross-validation error using the -cv flag in the ADMIXTURE command line. The analysis revealed the lowest cross-validation error when K was set to 11 (Supplementary Figure 2).

To uncover ancient splits and genetic relationships among populations, TreeMix v1.13 ^35^ was employed. The Önge genomes were utilised to establish the root of the maximum likelihood tree generated by TreeMix. Dataset 3 was used for plotting the maximum likelihood tree.

The statistical tool qpAdm ^36^ implemented in AdmixTools v5.1 ^17^ was employed to estimate ancestry proportions in the Koraga genomes originating from a mixture of reference populations by utilising shared genetic drift with a set of outgroup populations. Dataset 2, comprised of ancient and modern genomes, was employed for qpAdm and qpWave analysis. In the qpAdm analysis, the designation ‘Indus Periphery’ was applied to four ancient samples, specifically Rākhīgaḍhī_BA_Harappan, Shahr-i-Soktha_MLBA2, Shahr-i-Soktha_MLBA3 and Gonur2_BA ^16,27^. The Koraga were modeled as a combination of three source populations namely: Önge, Indus Periphery and a third population from Eurasia and Africa. The Mbuti from Africa, Scandinavian hunter-gatherers (SHG), Eastern hunter-gatherers (EHG), the Neolithic samples from Ganj Dareh in what today is western Iran, Neolithic Anatolians, Neolithic western Siberians, Hàn Chinese and the Karitiana from Brazil were used as the ‘Right’ outgroup populations (O8).

### Determination of the time of admixture

ALDER v.1.02 ^37^ was used to compute a weighted linkage disequilibrium (LD) analysis to infer the likely date of last admixture between the Koraga and other populations, considering a generation time of 30 years. The Koraga were modelled as the ‘admixpop’ (admixed population), and the remaining South Asian populations were considered as the ‘refpops’ (reference populations).

### Estimation of founder age using ASCEND

ASCEND v10.1.1 was used to estimate the founder age and intensity of the founder event in the Koraga population. The analysis was performed using two datasets for reliability. The first dataset included 14,539 SNPs, and the second dataset included 5,35,017 SNPs. Yoruba samples were considered as the outgroup. The founder age was calculated assuming a generation time of 25 and 30 years.

## Declarations and statements

### Ethics Approval

This study was performed in line with the principles of the Declaration of Helsinki. Approval was granted by Yenepoya Ethical Committee 1 (YEC-1/2021/052), affiliated with the Yenepoya deemed-to-be University.

## Author Contributions

JJS, RD and GvD contributed in conceptualisation. RD, MSM and SK collected the samples and performed data curation. JJS and RD performed statistical analysis and wrote the first draft of the manuscript. GvD reviewed, revised and redacted the entire manuscript draft. RD and MSM reviewed the final manuscript. All authors contributed to the manuscript and approved the submitted version.

## Acknowledgments

The authors acknowledge the participants, lab technicians and research staff involved in the study.

## Data Availability Statement

The raw genotype data of the participants cannot be shared for ethical reasons. However, the secondary data in the form of analysed files are available with the corresponding author and will be shared upon request.

