## Supplementary_File for "Novel 4,400-year-old ancestral component in a tribe speaking a Dravidian language"

Mohammed Shafiul Mustak

George van Driem

**Supplementary file**

As per the 2011 census, over 700 tribal populations, collectively known as Scheduled Tribes (STs), comprise 8.6% of the total population. Each tribal group is unique in terms of language, lifestyle, and social customs. Approximately 10% of the total tribal populations reside in Karnataka, a state in southern India. Notably, among these tribal communities, the Jenu Kuruba and Koraga are specifically classified as particularly vulnerable tribal groups (PVTGs).

There are various hypotheses about the Koraga people's origins, including local folklore and theories proposed by anthropologists, linguists, and historians. One of the most famous beliefs is that the Koragas are descendants of the "Chandala" caste, which refers to children born to a higher-caste woman and a lower-caste man. History also records a tribal leader named 'Habashika' who was exiled to the forests after being defeated by the Kadamba king (Thurston and Rangachari 1909). According to regional folklore, Hubbashika, a Koraga chief, purportedly governed the western coast of the southern Indian states of Karnataka and Kerala for 12 years during the 15th century CE (Chaudhuri and Bandopadhyay 2004) his army included people from the Koraga tribe. However, as per an ancient lore Hubbashika is said to have agreed to marry Kanayathi, the daughter of Lokaditya, a princess from the royal Varma family belonging to the Kadamba dynasty, that ruled parts of Karnataka. Allegedly, at the time of wedding, Hubbashika and his party were killed by Lokaditya's soldiers, and his loyalists that included the Koragas were expelled to the forest. The Koragas are believed to have surrendered to the Kadambas in lieu of protection and support but were purportedly marginalized by the victor troops. However, despite these stories, the genetic makeup of the Koraga tribe remains largely unknown, leaving their ancestral origin a mystery.

The Koraga people are classified into three endogamous groups: Soppu/Thappu (Leaves) Koraga, Kuntu (Cloth) Koraga, and Onti Koraga, according to Thurston and Rangachari's classification in 1909 (Thurston and Rangachari 1909).

**Supplementary Table 1.** Datasets used in this study

| **Dataset** | **Individuals** | **SNPs** | **LD pruned** | **Analysis** |
| --- | --- | --- | --- | --- |
| Dataset 1 | 1279 | 5,53,102 | NO | **smartpca**  We performed PCA using smartpca from the EIGENSOFT package (v18140)  lsqproject: YES  **ALDER v1.03**  mindis: 0.005  535017 SNPs used  **ASCEND v10.1.1**  Default settings  **Plink1.9**  --pca  **ADMIXTURE v1.3**  -cv |
| Dataset 2 | 4605 (Modern and ancient) | 14,539 | NO | **AdmixTools v5.1**  qpAdm and qpWave |
|  |  | 12,379 | YES  (--indep-pairwise 50 10 0.1) | **Admixtools 2** (find_graphs based on f2_from_precomp)  f3-statistics using Yoruba and Hàn as outgroups |
| Dataset 3 | 722 (Modern) | 14,539 | NO | **TreeMix v1.13** ML tree |
| Dataset 4 | 313 (GenomeAsia100K: Koraga related) | 190515 | NO  YES  (--indep-pairwise 50 10 0.1) | **Fst analysis using** dartR package:  gl.fst.pop  **Admixtools 2** (find_graphs based on f2_from_precomp) |

**PCA using Plink v1.9**

We performed a separate PCA using a different software on South Asian genomes for consistency. Dataset 1 was used for this purpose. PCA was performed using --pca command implemented in PLINK v1.9. Top two PCs were plotted using GraphPad Prism v10 (GraphPad Software, Boston, Massachusetts USA, [www.graphpad.com](http://www.graphpad.com)).

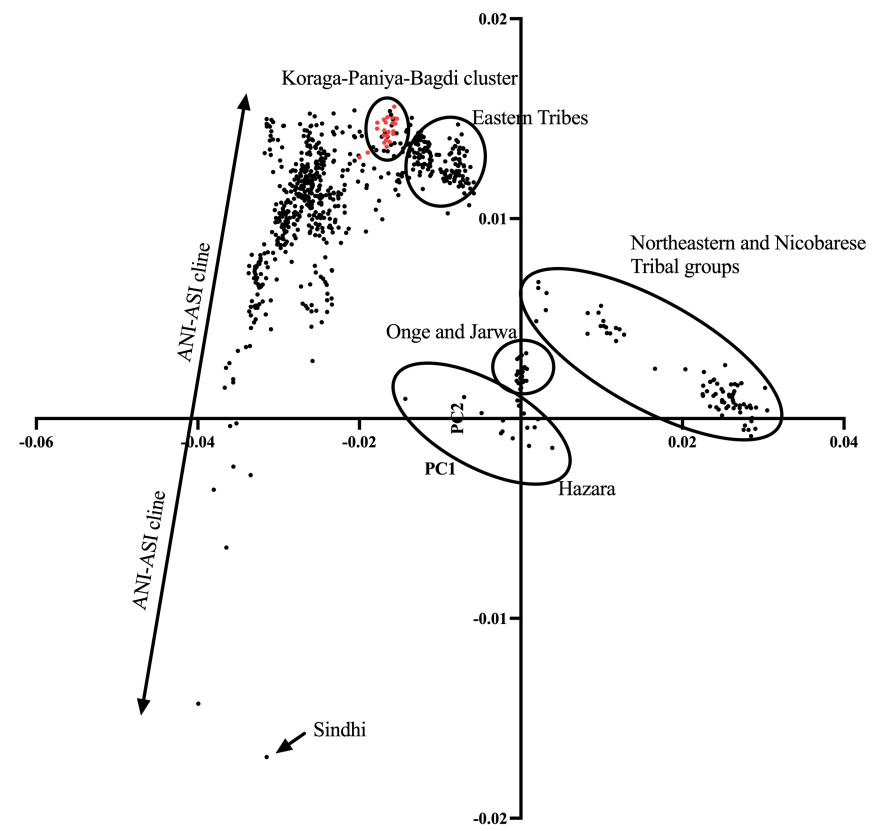

**Supplementary Fig. 1** Principal Component Analysis (PCA) using samples from Dataset 1. Only South Asian genomes were plotted to illustrate the genetic differentiation among the South Asian genomes. The X-axis (PC1) explained 28.19% variance while the Y-axis (PC2) explained 12.03% variance of the data. Notable populations are marked with circles. PCA was performed in PLINK v1.9. Top two PCs (PC1 and PC2), explaining the highest variance of the data were plotted in GrapPad Prism v10.

The Principal Component Analysis (PCA) using Dataset 1 (see Methods) revealed a previously described Ancestral North Indian (ANI)–Ancestral South Indian (ASI)–Ancestral Austroasiatic (AAA) cline along the vertical principal component (PC2) (Basu et al. 2016; Das and Upadhyai 2018) with Sindhi, Kalash and other Pakistani populations, clustering at one extreme of the cline, and the indigenous groups and tribal populations on the other end of the same (Supplementary Fig. 1). Consistent with the previous studies, ASI-AAA-Ancestral Tibeto-Burman (ATB) cline was observed along the horizontal principal component (PC1) with the tribal populations from the mainland clustering at one end, while the populations from the Northeast and Nicobar Islands of India clustered at the other end (Supplementary Fig. 1). The Koraga samples constituted a unique cluster, distinct from both the ANI-ASI cline and independent of the Eastern tribe cluster. Additionally, the Koraga genomes clustered together with primitive Dravidian speaking Indian populations, such as Paniya and Bagdi. This suggests an early divergence of Koraga genomes from the upper caste Dravidian speaking populations in southern India (Supplementary Fig. 1). These PCA results are in congruence with smartpca results (Fig. 1 main text).

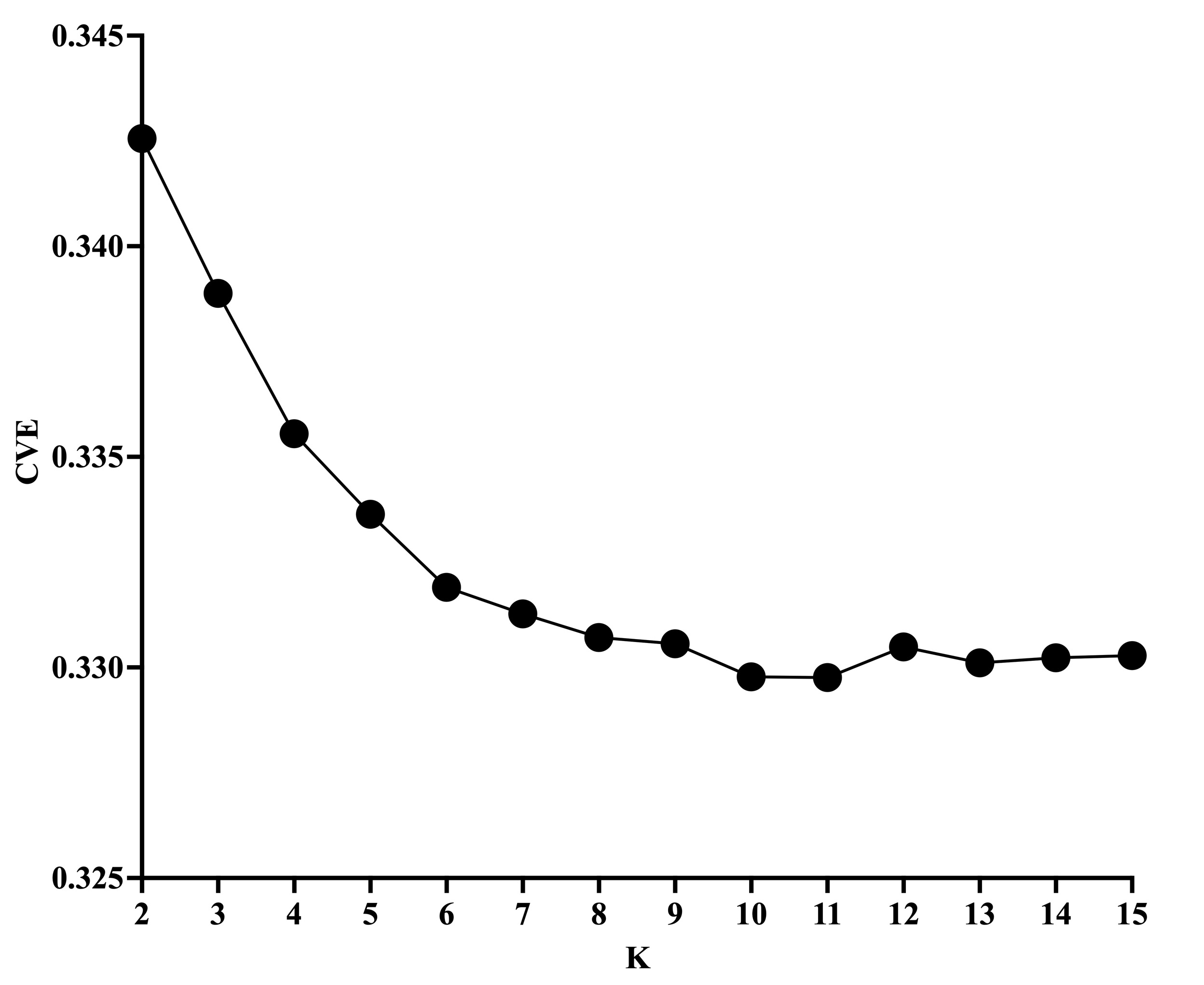

**Supplementary Fig. 2** Cross-validation error (CVE) for K values in the ADMIXTURE analysis. The lowest cross-validation error (CVE) was observed for K = 11.

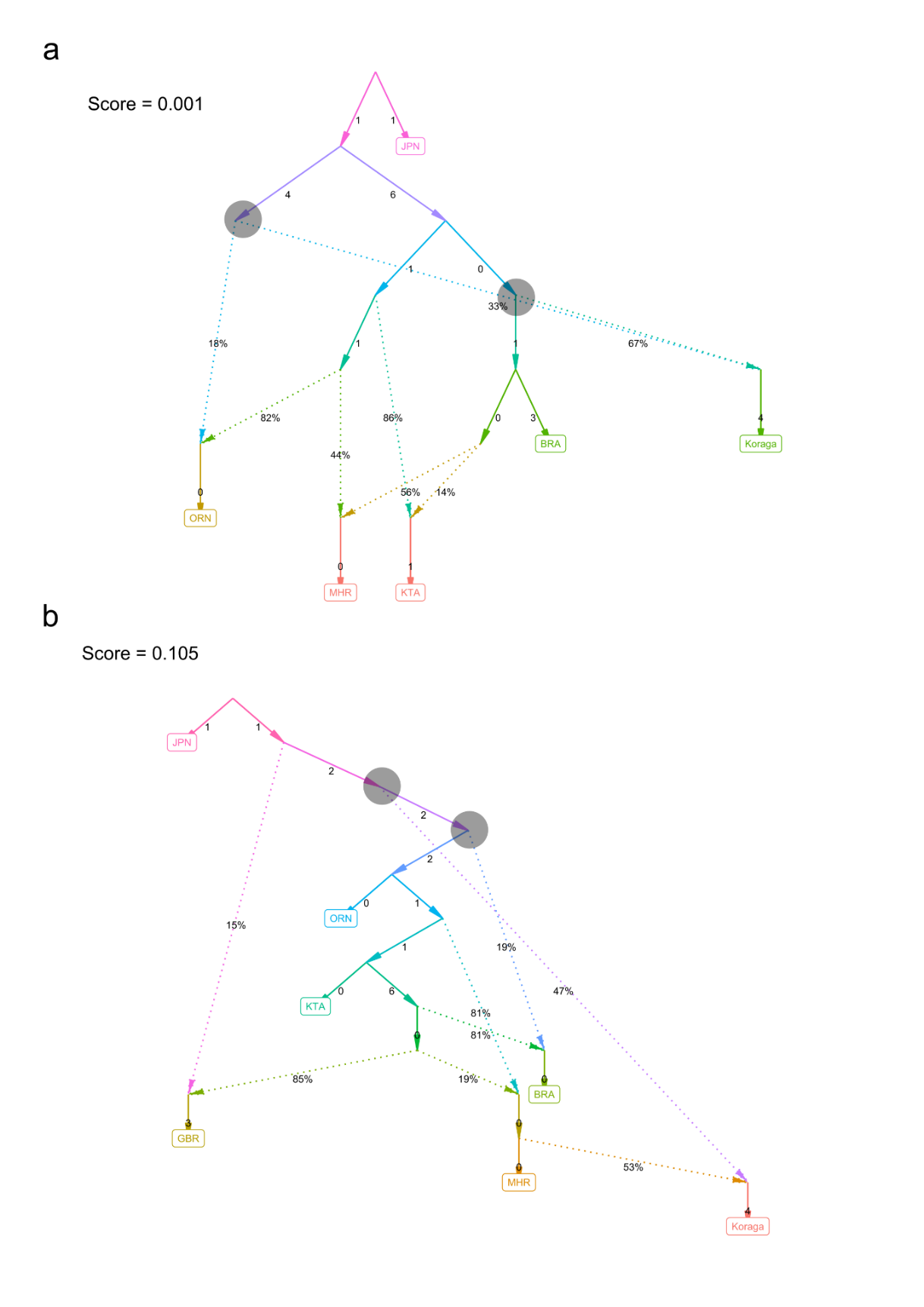

**Supplementary Fig. 3** Admixture graphs showing admixture possibilities in the present day populations. JPN is used as an outgroup. (a) Koraga shares its ancestry with Brahui and Oraon populations. Mahar and Kota tribes derive their ancestry from the ancestors of Brahui and Oraon. (b) Koraga shares ancestry with the ancestor of Mahar and from the ancestral lineage of Brahui, Oraon and Kota. Kota is a Dravidian speaking south Indian tribe. Mahar is an Indo European language speaking Western tribe, Oraon is a North Dravidian speaking Eastern tribe. Brahui is a North Dravidian speaking tribe found in Balochistan, Pakistan.

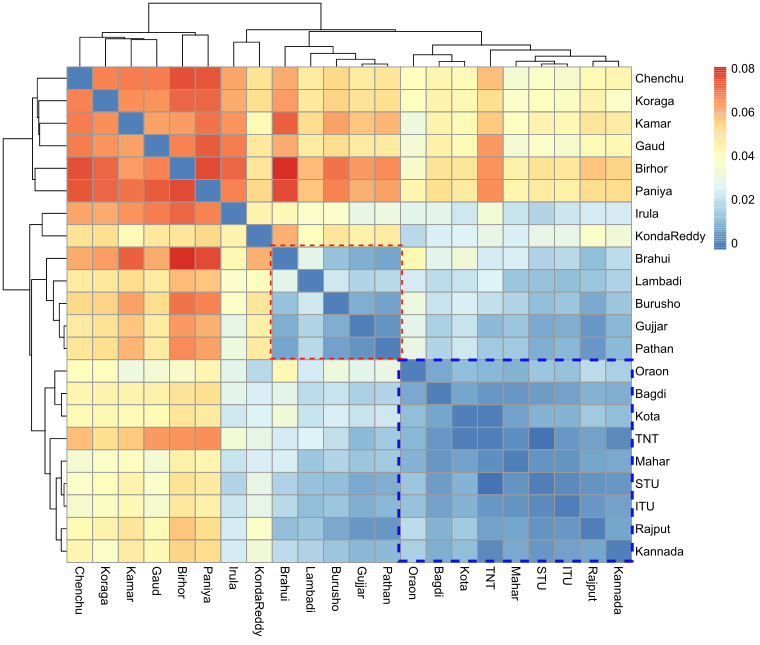

**Supplementary Fig. 4** Fst heatmap of selected populations from the Indian subcontinent. Blue square represents a cluster that includes Oraon, Bagdi, Tanti (TNT) and Mahar populations with successful ALDER LD decay with Brahui and Koraga. Other populations in this cluster include Tamil (STU), Teludu (ITU), Rajput, Kota and Kannada which are found on the Southern, Western and North Western India. Red square includes populations from the North Western region with higher West Eurasian component. Koraga clusters with Gaud, Birhor, Paniya, Chenchu and Kamar tribes that are found as outliers in the PCA plot.

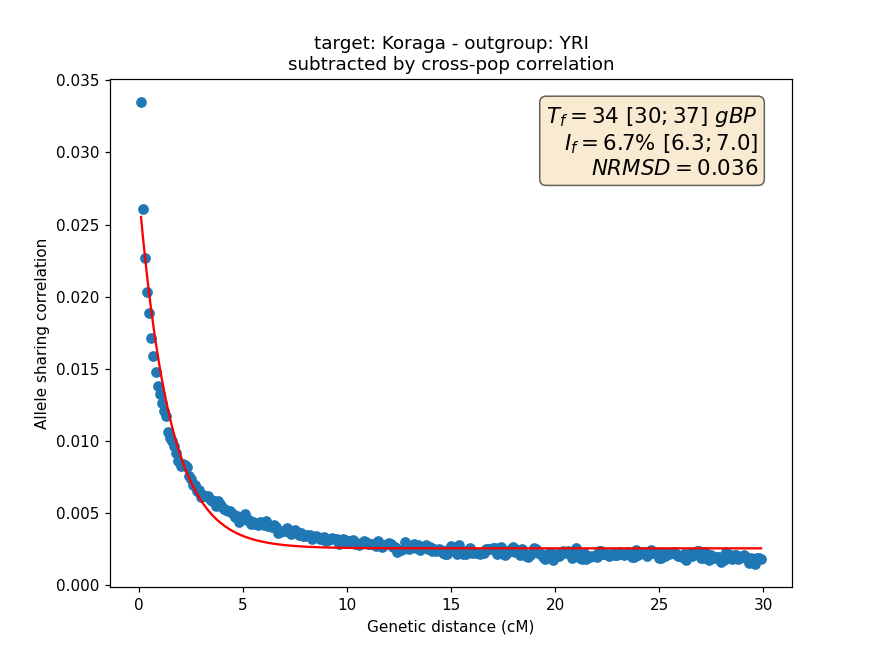

**Supplementary Fig. 5** Results of ASCEND run for Koraga. YRI was considered as an outgroup. Founder age estimate and intersity are listed in Supplementary Table 2.

**Supplementary Table 2** Founder age estimate and founder event intensity in Koraga tribe

| Dataset | Founder Age Estimate | | | Founder Intensity (%) |
| --- | --- | --- | --- | --- |
|  | Generations before sampling (95% CI) | 30 years/generation | 25 years/generation |  |
| 14539 SNPs | 30 (22;37) | 900 (660-1110) | 750 (550-925) | 8.2 (7.4;9.1) |
| 535017 SNPs | 34 (30;37) | 1020 (900-1110) | 850 (750-925) | 6.7 (6.3;7.0) |
